# Diverse noncoding mutations contribute to deregulation of cis-regulatory landscape in pediatric cancers

**DOI:** 10.1101/843102

**Authors:** Bing He, Peng Gao, Yang-Yang Ding, Chia-Hui Chen, Gregory Chen, Changya Chen, Hannah Kim, Sarah K. Tasian, Stephen P. Hunger, Kai Tan

**Affiliations:** Division of Oncology and Center for Childhood Cancer Research, Children’s Hospital of Philadelphia, Philadelphia, PA 19104, USA; Department of Pediatrics, Perelman School of Medicine, University of Pennsylvania, Philadelphia, PA 19104, USA; Medical Scientist Training Program, University of Pennsylvania, Philadelphia, Pennsylvania 19104, USA; Department of Biomedical and Health Informatics, Children’s Hospital of Philadelphia, Philadelphia, Pennsylvania 19104, USA; Department of Genetics, Perelman School of Medicine, University of Pennsylvania, Philadelphia, PA 19104, USA; Department of Cell and Developmental Biology, Perelman School of Medicine, University of Pennsylvania, Philadelphia, PA 19104, USA; Abramson Cancer Center, Perelman School of Medicine, University of Pennsylvania, Philadelphia, PA 19104, USA

## Abstract

Interpreting the function of noncoding mutations in cancer genomes remains a major challenge. Here we developed a computational framework to identify risk noncoding mutations of all classes by joint analysis of mutation and gene expression data. We identified thousands of SNVs/small indels and structural variants as candidate risk mutations in five major pediatric cancers. We experimentally validated the oncogenic role of *CHD4* overexpression via enhancer hijacking in B-ALL. We observed a general exclusivity of coding and noncoding mutations affecting the same genes and pathways. We showed that integrated mutation signatures can help define novel patient subtypes with different clinical outcomes. Our study introduces a general strategy to systematically identify and characterize the full spectrum of noncoding mutations in cancers.

## Introduction

Extensive research efforts have identified recurrent mutated genes in multiple types of childhood cancer including several recent pan-cancer studies which have defined landscapes of coding mutations in common pediatric cancers (1-4). However, current cancer mutation landscapes are far from complete without systematic analysis of the noncoding portion of the genome. For instance, recurrent leukemia-associated genetic alterations cannot be identified in 10%∼20% of children with acute leukemias (1,3-5), thereby making it challenging to design targeted therapies for such patients.

The vast majority of somatic mutations in cancer genomes occur in noncoding regions because 98% of the human genome is made up of noncoding sequences, and the somatic mutation rate of noncoding regions is similar to that of coding regions (1). The Therapeutically Applicable Research To Generate Effective Treatments (TARGET) project has sequenced over 1000 genomes from five common pediatric cancers. Similarly, The Cancer Genome Atlas (TCGA) project has molecularly characterized over 20,000 primary cancer genomes spanning 33 types of adult cancers. Despite this rapid accumulation of whole genome sequences (WGS) for both pediatric and adult cancers, identification and interpretation of the functional impact of noncoding mutations remains challenging.

Noncoding regulatory sequences, particularly enhancers and promoters, are key determinants of tissue-specific gene expression. Multiple mutation types have been reported to disrupt enhancers and promoters and expression of their target genes, including single nucleotide variants (SNVs), small insertion and deletions (indels), and large structural variants (SVs), including deletions, insertions, duplications, inversions, and translocations. Most prior studies of noncoding mutations have focused on SNVs and small indels and revealed a number of noncoding causal mutations. One example is the SNP rs2168101 G>T located in the enhancer of *LMO1*, which disrupts GATA3 binding and *LMO1* expression in neuroblastoma patients (6). Another example is the heterozygous indels (2-18 bp) located at −7.5 kb from *TAL1* transcription start site in T-acute lymphoblastic leukemia (T-ALL), which introduces new binding sites for MYB to create a super-enhancer upstream of *TAL1* (7).

Only about 30% of SVs result in in-frame gene fusions that join the protein coding regions of two genes (8). The majority of SV break points are located in noncoding regions and do not change gene structure. Although less well studied compared to SNVs and small indels, several seminal studies have revealed oncogenic roles of noncoding SVs by redirecting enhancers/promoters to oncogenes or from tumor suppressor genes (9,10). Such enhancer rearrangement events have been identified in diverse cancer types, suggesting that this could be a common mechanism of oncogenesis. For instance, interstitial deletion in the pseudoautosomal region 1 (PAR1) of the X/Y chromosomes places the transcription of *CRLF2* (cytokine receptor-like factor 2) under the control of the *P2RY8* (purinergic 2 receptor Y 8) enhancer in Philadelphia chromosome-like and Down Syndrome-associated B-ALL (11). Other examples include complex structural variants in childhood medulloblastoma, which rearranges the *DDX31* (DEAD-Box Helicase 31) enhancer to *GFI1B* (growth factor independent 1B) (9), and t(3;8) in B-cell lymphoma that rearranges the *BCL6* enhancer to *MYC* (10).

Given the prevalence of noncoding mutations and the drastic increase of whole genome sequencing data, novel computational methods are critically needed to systematically identify risk noncoding mutations. Here, we introduce the PANGEA method (<u>P</u>redictive <u>A</u>nalysis of <u>N</u>oncoding <u>G</u>enomic <u>E</u>nhancer/promoter <u>A</u>lterations), a general computational framework for systematic analysis of noncoding mutations and their impact on gene expression. This method simultaneously identifies all classes of somatic mutations that are associated with gene expression changes, including SNVs, small indels, copy number variations, and structural variants. Using PANGEA, we have conducted a pan-cancer analysis of noncoding mutations in 501 pediatric cancer patients of five histotypes with matched WGS and RNA-Sequencing (RNA-Seq) data generated by the TARGET project. We identified a comprehensive list of recurrent noncoding mutations as candidate risk mutations in these cancers. An integrated analysis of both coding and noncoding mutations revealed distinct pathways affected by either coding and noncoding mutations. We also show that integrated mutation signatures can help define novel patient subtypes with different clinical outcomes.

## Results

### A comprehensive catalog of recurrent noncoding mutations

The TARGET data portal (https://ocg.cancer.gov/programs/target/data-matrix) contains matched WGS and RNA-Seq data for 501 patients, including 163 patients with B-cell acute lymphoblastic leukemia (B-ALL), 153 patients with acute myeloid leukemia (AML), 100 with neuroblastoma (NBL), 53 with Wilms tumor (WT), and 32 with osteosarcoma (OS). All patients have WGS data for both tumor and germline or remission samples, which defined alterations as germline or somatic.

We identified somatic SNVs and small indels (< 60bp) for the five cancer types using tumor and remission samples (Figure S1A). We evaluated the accuracy of our mutation calling pipeline using two approaches. First, we used a benchmark set generated by Zook et al. (12). It consists of a set of high-confidence SNVs and indels in subjects NA12878 and NA24631 from the 1000 Genome Project, identified by integrating 14 data sets using five sequencing technologies, seven read mappers and three variant callers. Using this benchmark set, we estimated the precision of our pipeline to be ∼95% (Table S1). Second, the TARGET project provides a list of high-confidence SNV calls in B-ALL, AML, NBL, and WT that were validated using multiple experimental protocols including whole exome sequencing (WES), RNA-Seq and targeted sequencing. Of the 735 high-confidence calls, our analysis pipeline identified 681 (92%) of them, suggesting high sensitivity of our pipeline (Figure S1B).

Across the five cancer types, the average mutation rate ranges from 0.16 SNV/indel per million bases (Mb) in AML to 0.55 per Mb in OS (Figure S1C). The higher somatic mutation rate in OS is consistent with a previous WGS study, in which regional clusters of hypermutation, termed kataegis, were identified in 85% of pediatric OS patients (13). As expected, over 98% of the identified mutations are located in noncoding regions, while less than 1% of the mutations are located in coding regions (Figure S1D).

Next, we used Delly2 (14) and Lumpy to identify structural variants (SVs), including large deletions (DELs), tandem duplications (DUPs), inversions (INVs), and translocations (TRANs). We only kept SVs that were called by both methods and passed additional filtering criteria (Figure S2A) as the final set of identified SVs. In total, we identified 26,757 SVs across 501 patients. The average number of identified SVs per patient ranges from 18 in WT to 71 in OS (Figure S2B). These numbers are comparable with published data by the International Cancer Genome Consortium (ICGC) and The Cancer Genome Atlas (TCGA) consortium. Previously, Roberts *et al*. identified and experimentally validated 21 SVs that joined exons of two genes in frame in B-ALL patients (15). We identified 20 of those SVs, suggesting high sensitivity of our pipeline. Among the identified SVs, 9,831 are in-frame changes and potentially generate fusion genes. We evaluated the fusion gene predictions by comparing to matched RNA-Seq data from the patients. Of the 9,831 fusion genes predicted by our pipeline, 2,323 of them were supported by RNA-Seq data from the same patients. The TARGET consortium experimentally tested 12 known translocations in 212 leukemia patients. Based on this set of validated SVs, our SV calling pipeline has an accuracy of 98% (Figure S2C, Table S2). Taken together, using multiple benchmarking data sets, we show that our SV calling pipeline is highly accurate.

### Noncoding mutations disrupting enhancer/promoter sequences or enhancer-promoter interactions

The functional consequence of noncoding mutations is difficult to interpret without comprehensive annotation of noncoding regulatory DNA sequences in cancer genomes. To this end, we used publicly available epigenomic data of disease-relevant cell types to construct an enhancer catalog for the five cancer types in this study (see Methods). Specifically, we used ChIP-Seq data of histone modification marks (H3K4Me1, H3K4Me3, H3K27Ac) to predict transcription enhancers using the chromatin signature identification by artificial neural network (CSI-ANN) algorithm (Table S3) (16). In total, we identified 282,021 enhancers for the five cancer types at the False Discovery Rate (FDR) of 0.05 (Figure S3A). Overall, 91% of predicted enhancers are supported either by ATAC-Seq data in cancer-relevant cell types or by sequence conservation across 20 mammalian genomes (Figure S3A) or both, suggesting high quality of the predicted enhancers. Next, we predicted target gene(s) of each enhancer using the Integrated Method for Predicting Enhancer Targets (IM-PET) algorithm (17), using public histone modification ChIP-Seq and RNA-Seq data in disease-relevant cell types. In total, we predicted 635,096 enhancer-promoter (EP) pairs across the five cancer types (Figure S3A). We compared the predicted EP pairs with published high-resolution Hi-C/ChIA-PET data in human B cells, myeloid cells and kidney cells (Table S3, Methods). Seventy-four percent (317,698) of our EP predictions are supported by either Hi-C or ChIA-PET data (Figure S3B), suggesting high quality of our predictions.

To identify recurrent mutations that disrupt either enhancer/promoter sequences or enhancer-promoter interactions, we intersected the catalog of somatic mutations with the catalogs of enhancers/promoters and EP interactions. Across the five cancer types, 16% SNVs or indels overlap with enhancers/promoters in cell types relevant to a given pediatric cancer. In comparison, only 8% of all SNVs across 34 TCGA cancer types (as a control set) overlap with enhancers/promoters in the relevant cell types (average p-value = 5e-13) (Figure S4A). The identified CNVs also have significantly higher overlap with the enhancers in relevant cell types compared to all CNVs reported by the TCGA consortium (15% vs 7%, average p-value = 2e-11) (Figure S4B). The break points of inversions and translocations are significantly enriched between predicted EP pairs compared to all break points reported by the TCGA consortium (45% vs 36%, average p-value = 1e-18) (Figure S4C). Taken together, these results suggest that various types of somatic mutations can disrupt the cis-regulatory landscapes of pediatric cancers, especially cis-regulatory elements involved in regulating tissue-specific gene regulation of the specific tumor histotype.

### Systematic identification of candidate risk noncoding mutations

To help prioritize noncoding mutations, we developed the PANGEA method to systematically identify recurrent noncoding mutations that disrupt the transcriptional regulation of a gene. Unlike previous methods, PANGEA uniquely considers all types of noncoding mutations, including SNVs, small indels, CNVs, and SVs. These mutation types can either disrupt enhancer/promoter sequences or disrupt the interactions between enhancers and promoters which is critical for transcription activation. After tabulating all such mutations, we use weighted elastic net to perform a regression analysis of gene expression on the sets of noncoding mutations across the patient cohort (Figure 1A). Because the EP interactions are predicted computationally, we use EP prediction scores as the weights for each predictor (mutation) in order to include a confidence measure of EP interactions in our regression model. Candidate risk mutations are predicted based on the statistical significance of the corresponding regression coefficients (see Methods for details). The PANGEA software package is available at https://github.com/tanlabcode/PANGEA.

**Figure 1.**
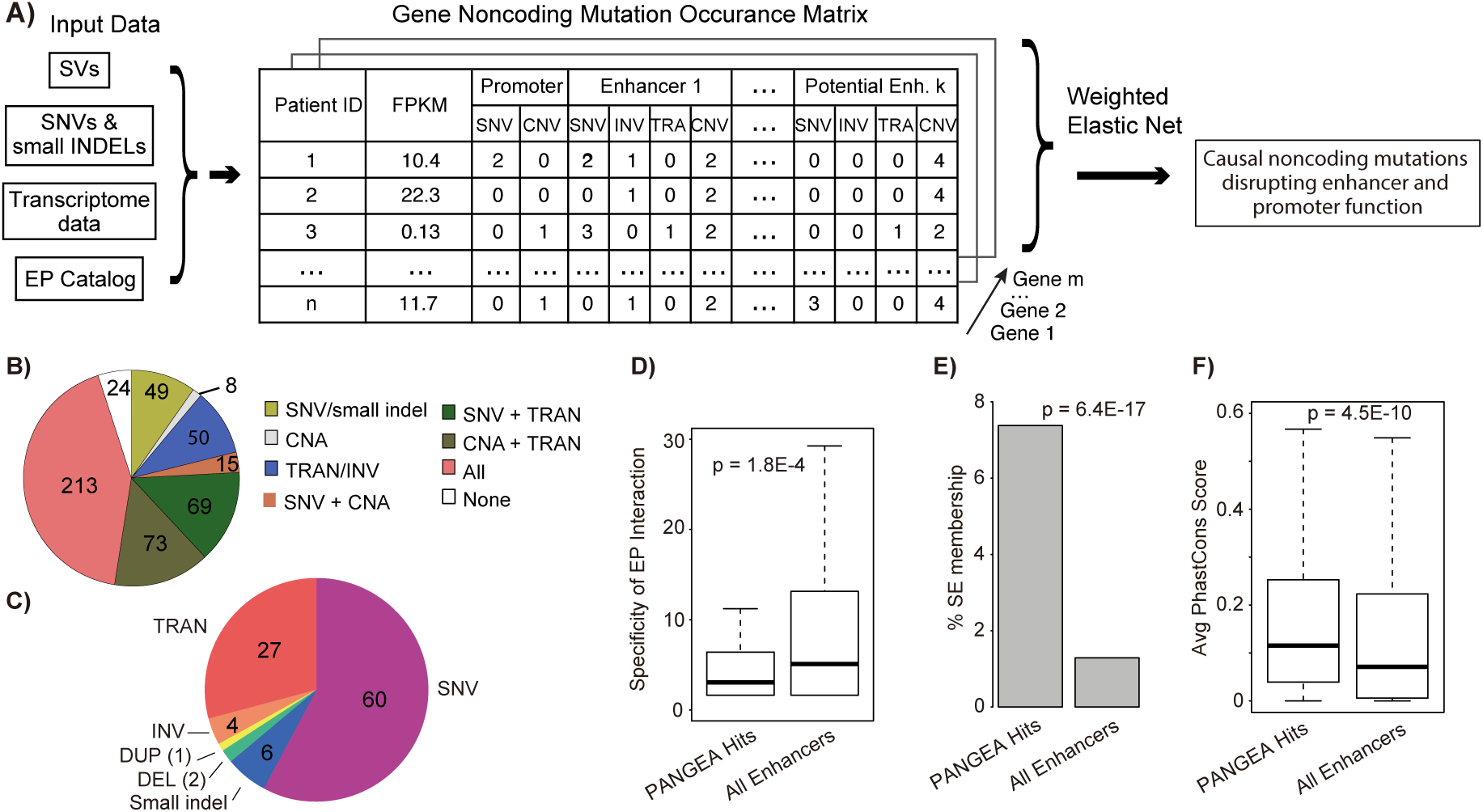
The PANGEA algorithm for simultaneous identification of candidate risk noncoding mutations of all classes. **A)** Overview of the PANGEA algorithm. Given a catalog of enhancer-promoter (EP) interactions, a catalog of somatic mutations, and transcriptome data, the algorithm first constructs a count matrix of various classes of mutations disrupting the function of enhancer(s)/promoter(s) of each gene, including small mutations and structural variants. Weighted elastic net regression is then used to identify mutations that are significantly correlated with changes in target gene expression. **B**) Fraction of patients stratified by different classes of noncoding mutations affecting enhancers/promoters. **C)** Proportion of genes whose expression is disrupted by different classes of predicted risk noncoding mutations. SNV, single nucleotide variant; DEL, deletion; DUP, duplication; TRAN, translocation. **D)** Mutations predicted by PANGEA affect enhancers that have higher tissue specificity. Enhancer specificity is measured by the number of cell types in which the enhancer is observed to be active. P-value of one-sided t-test is shown (n=282,021); **E)** Mutations predicted by PANGEA affect higher percentage of super enhancers. p-value of hypergeometric test is shown (n=282,021); **F)** Mutations predicted by PANGEA affect enhancers with higher sequence conservation across 20 mammalian species. Enhancer conservation is measured by the average PhastCons score of the enhancer sequence. P-value of one-sided t-test is shown (n=282,021).

Using multiple-testing adjusted p-value < 0.05 as the cutoff, we identified 1,405 genes whose expression changes can be predicted by SNVs/small indels in their enhancers/promoters, 55 genes whose expression changes can be predicted by CNVs in their enhancers, 1,082 genes whose expression changes can be predicted by SVs that disrupt their EP interactions (Table S4). In total, disruption of enhancer function by recurrent noncoding mutations were found in 477 out of 501 pediatric cancer patients (95%) (Figure 1B). The quantile–quantile plot shows large deviation of the observed P values for the predicted risk noncoding mutations compared to those of random expectations using an independent ICGC cohort (n=2715 donors), suggesting low false prediction rate (Figure S5, Supplementary Methods). Over half of the predicted risk noncoding mutations are SNVs and small indels (66%), followed by translocations (27%) and other types of SVs (Figure 1C). However, when adjusted by the overall frequency of each type of mutation, SVs become the most frequent type of risk noncoding mutations (Figure S6).

The enhancers that are affected by risk noncoding mutations are more cell-type-specific compared to all enhancers in our enhancer catalog (Figure 1D). In addition, these enhancers have more overlap with published super enhancers in cell types that are relevant to the given cancer type (18) (Figure 1E). Finally, these enhancers have high levels of sequence conservation across 20 mammalian species (Figure 1F). Taken together, these results provide additional support to the predicted risk noncoding mutations.

### Risk enhancer rearrangements

Several seminal studies have reported ‘enhancer hijacking’, also known as oncogenic rearrangement of enhancers due to translocation/inversion (9,10,19-22). In this study, we performed a systematic analysis of enhancer hijacking events across the five cancer types. Amongst 501 total patients, 405 patients (81%) have at least one predicted enhancer hijacking event in their genomes. We identified several known enhancer hijacking events. For example, we found the t(14;X)(q32;p22) translocation in 12 B-ALL patients, which hijacks multiple enhancers of *IGHV* to the vicinity of *CRLF2* resulting in significant overexpression of *CRLF2* in these patients (Figure S7A). We also discovered enhancer hijacking from three different genomic loci to the common *TERT* gene locus (t(10;5)(p22;p15), t(5;5)(q34;p15), and t(5;5)(q12;p15) (Figure S7B). These *TERT*-related enhancer hijacking events were observed in 15 neuroblastoma patients and the resultant translocations led to significant overexpression of *TERT* in these patients (Figure S7B).

However, the majority of our predicted enhancer hijacking events have not been previously reported. One such event involves hijacking of enhancers to upregulate *CHD4*, which has three translocation partners (t(12;22)(p13;q13), t(12;19)(p13;p13), t(9;12)(p24;p13)). In total, 12 B-ALL patients had this translocation. Most translocations in this region result in fusion genes involving a nearby gene *ZNF384*. A previous study has focused on the function of *ZNF384* fusion genes (23). However, in the TARGET cohort, we observed 2 patients in which the translocation break point is actually located downstream of *ZNF384* and did not generate an *ZNF384* fusion (Figure 2A). Furthermore, ZNF384 fusion is not correlated with patient prognosis. Together, these data suggests a driver of oncogenesis that is independent of *ZNF384*. In all patients with the translocation, we found that enhancers such as that of *EP300, TCF3*, and *SMARCA2* are hijacked to chr12.p13. The expression level of *CHD4*, a gene adjacent to *ZNF384*, is significantly increased in these patients (Figure 2B). Similarly, in a recent published mixed phenotype acute leukemia (MPAL) cohort, we also observed *CHD4* expression increase in the patients with *ZNF384*-involved rearrangement (3) (Figure S8). Moreover, B-ALL patients with the enhancer hijacking event have significantly shorter time to relapse (Figure 2C). We thus hypothesize that enhancer hijacking translocation events that bring potent enhancers to the promoter of *CHD4* may be an independent oncogenic driver in B-ALL.

**Figure 2.**
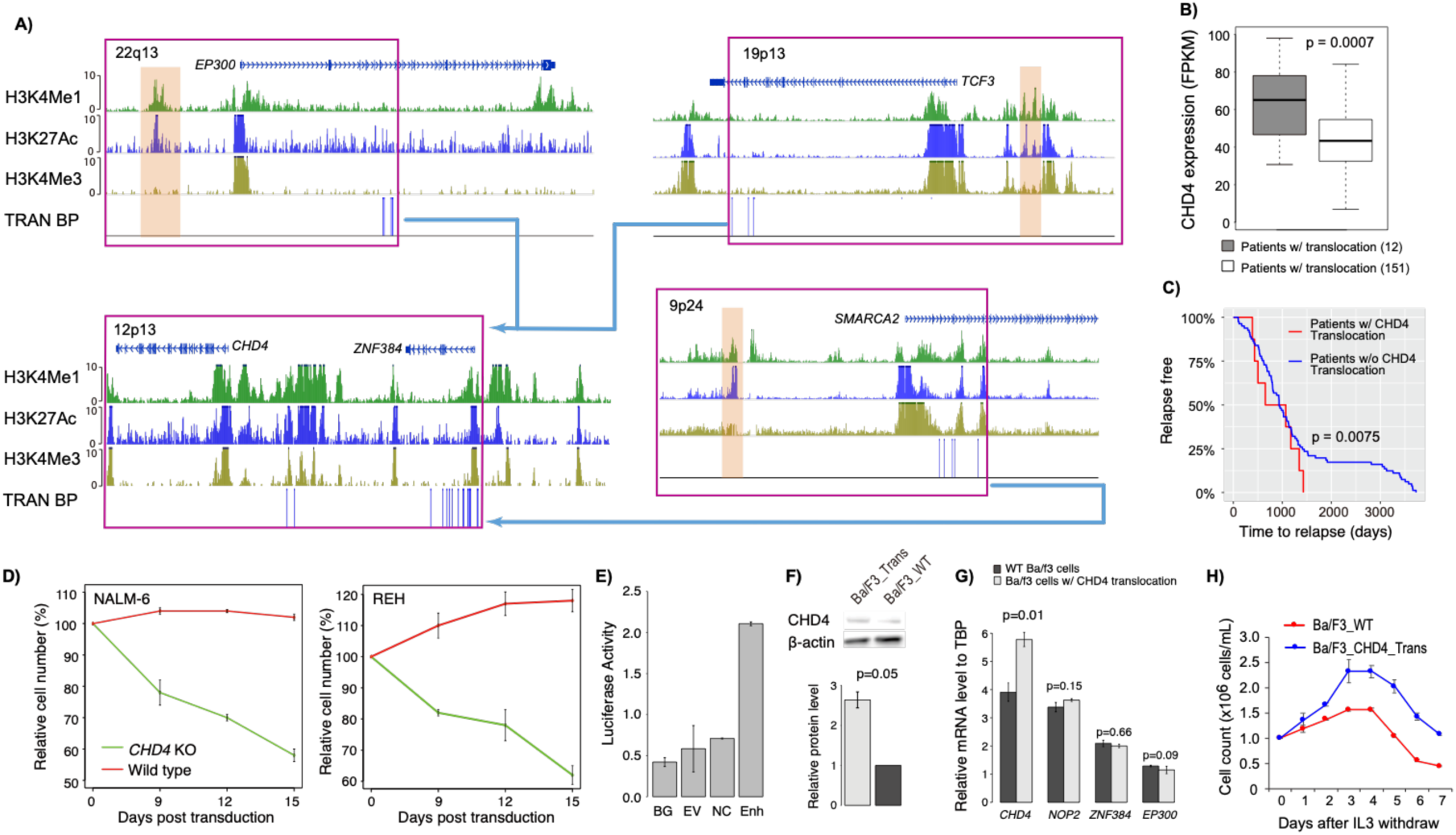
Enhancer hijacking to *CHD4* by translocation in B-ALL patients. **A)** Translocations result in enhancer hijacking to *CHD4* in B-ALL. Three different translocation partners are identified in the B-ALL patient cohort. Shown tracks are histone modification ChIP-Seq data of human CD19^+^ B cells (identifying enhancers), SNVs and SV break points (BPs). For break point track, each vertical line represents a patient. The hijacked enhancers predicted to regulate *CHD4* are highlighted in brown. **B)** Expression levels of *CHD4* in patients with and without translocations. P-value of one-sided t-test is shown (n=163). **C)** B-ALL patients with *CHD4* translocation have shorter time to relapse. P-value of log-rank test is shown (n=163). **D)** *CHD4* knockout impairs the growth of B-ALL cell lines, NAML-6 and REH. *CHD4* knockout was done using CRISPR-Cas9. **E)** Luciferase reporter assay of hijacked enhancer. BG, no DNA vector control; EV, vector containing no enhancer; NC, negative control, genomic region without enhancer histone mark H3K4me1 and H3K27ac; Enh, test *EP300* enhancer. **F)** Western blots of CHD4 protein level in Ba/F3 cells with and without introduced translocation. P-value of one-sided t-test is shown (n=2). **G)** Relative mRNA levels of *CHD4, NOP2, ZNF384*, and *EP300* in Ba/F3 cells with and without introduced translocations. P-values were calculated using one-sided t-test (n=2 for all tests). **H)** Murine Ba/F3 cells with induced translocation undergoes oncogenic transformation. Enhancer hijacking translocation was induced by CRISPR-Cas9. Error bars represent one standard deviation.

*CHD4* (chromodomain-helicase-DNA-binding protein 4) is a component of the nucleosome remodeling and deacetylase (NuRD) complex, which plays an important role in B-cell development by regulating B-cell-specific transcription (24). In addition, *CHD4* is known to function as a repressor of several tumor suppressor genes, and inhibition of *CHD4* reduces the growth of AML and colon cancer cells (25,26). These data suggest a role of *CHD4* in the oncogenesis of B-ALL. To investigate the potential role of *CHD4*, we first identified differential expressed genes in the patients with *CHD4* overexpression. In total, there are 666 up-regulated and 922 down-regulated genes in patients with *CHD4* rearrangements (Table S5). Using published TF ChIP-Seq data in human GM12878 cells (27), we found CHD4 binding sites are significantly enriched at enhancers or promoters of the down-regulated genes (Figure S9A). Among the 157 down-regulated genes with CHD4 binding sites, several of them encode well-known regulators of B cell development including *PAX5, IRF4, TCF3* and *EBF1* (Figure S9B, C). These results suggest a role of CHD4 in B-ALL through regulating the expression of key TFs in B-cell development.

To test the potential oncogenic role of *CHD4* in B-ALL, we knocked out *CHD4* in the NALM-6 and REH B-ALL cell lines. We performed growth competition assays to compare the growth phenotypes of *CHD4* knockout with non-knockout leukemic cells. Both NALM-6 and REH cells showed impaired growth with *CHD*4 knockout (Figure 2D). Next, we introduced the translocation t(6;15)(qF2;qE1) in murine Ba/f3 cells using CRISPR-Cas9. The translocation break points were designed to be located in the intergenic regions near *CHD4* and *EP300*, placing the *EP300* enhancer (Figure 2E) upstream of the *CHD4* promoter without creating a fusion gene (Figure S10A). In Ba/f3 cells with the translocation, we observed increased expression of *CHD4* at both mRNA and protein levels (Figure 2G, H). In contrast, expression of nearby genes was not increased in cells with the translocation (Figure 2G). More importantly, the introduced translocation enables Ba/f3 cells to proliferate in the absence of IL-3 (Figure 2H), suggesting oncogenic transformation. Additionally, the expression levels of *TCF3, PAX5*, and *EBF1* were significantly decreased in the cells with CHD4 rearrangement, suggesting the role of CHD4 in regulating those key regulators (Figure S10B). In summary, these results strongly support an oncogenic role of *CHD4* in B-ALL due to enhancer hijacking.

### Risk enhancer copy number alterations

We found that 309 patients in the TARGET cohort (62%) had enhancer amplification/deletion and associated target gene expression change. *MYCN* is frequently amplified in neuroblastoma patients (28). In the NBL cohort, we observed that the body of the *MYCN* gene is amplified in 34 patients. However, we also observed 11 patients with amplification of the *MYCN* enhancer rather than the gene body (Figure 3A). *MYCN* expression level is significantly higher in those patients compared to the patients without *MYCN* amplification (Figure 3B). Moreover, patients with only *MYCN* enhancer amplification have shorter time to relapse compared with other patients (Figure 3C), including patients with *MYCN* gene body amplification. This result suggests that enhancer amplification alone is sufficient to up-regulate *MYCN* and drive aggressive neuroblastoma. Another example involves deletion of the *ATG3* enhancer in 15 B-ALL patients (Figure S11A), resulting in decreased *ATG3* expression in those patients (Figure S11A). Autophagy related 3 (*ATG3*) is a ubiquitin-like-conjugating enzyme and plays a role in the regulation of autophagy; downregulation of *ATG3* has been reported in myelodysplastic syndrome (MDS) patients progressing to leukemia (29).

**Figure 3.**
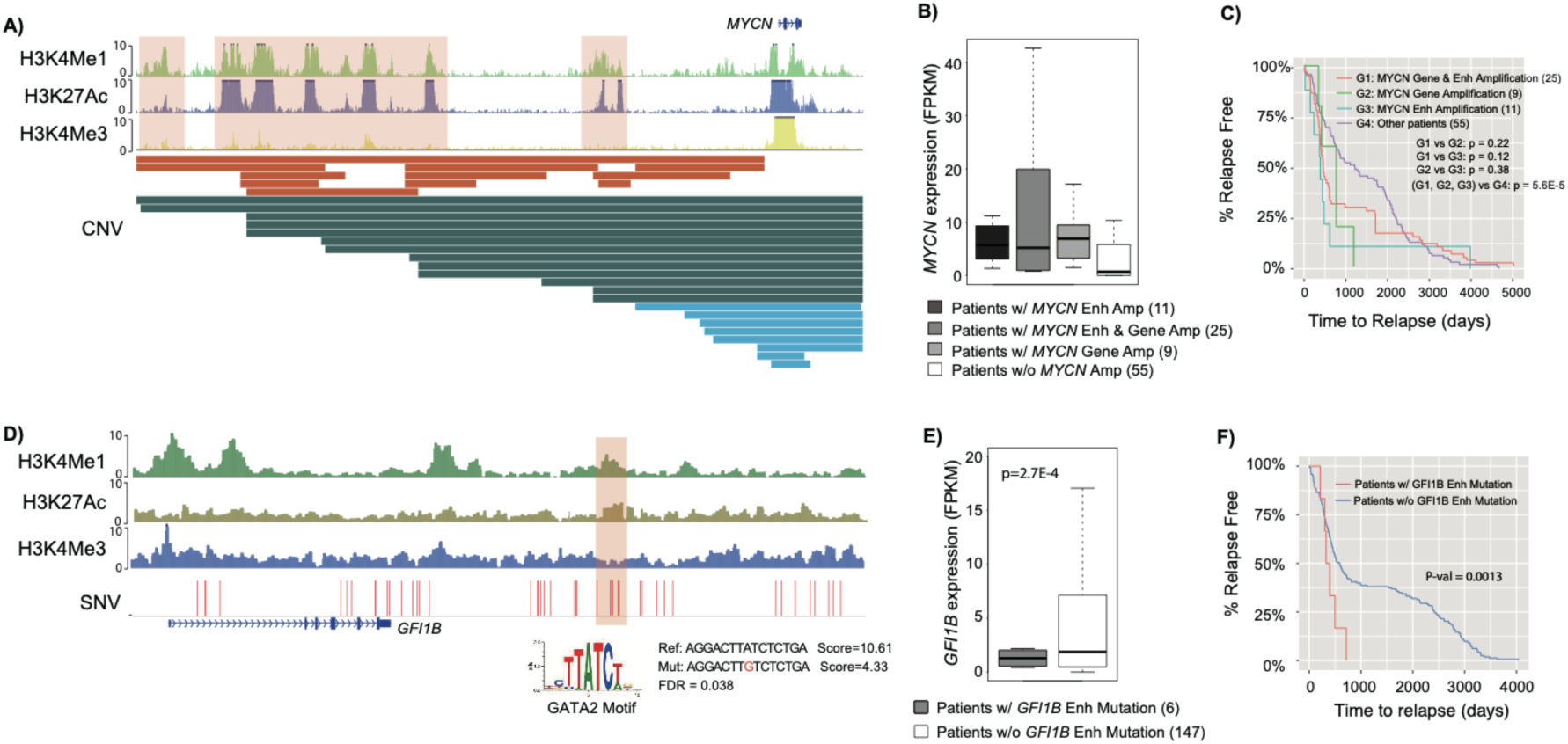
Examples of disruption of enhancer function by copy number alteration and point mutation. **A)** Segmental duplications result in amplification of *MYC* enhancer in neuroblastoma. Shown tracks are histone modification ChIP-Seq data in human neurocrest cells and identified CNVs. Color bars in CNV track indicate MYCN amplification types. Orange, amplification of MYCN enhancers; Green, amplification of both MYCN enhancers and gene body; Blue, amplification of MYCN gene body only. **B)** Expression level of *MYC* in patients with and without copy number change. **C)** Kaplan-Meier plots of time to relapse for NBL patients with *MYC* gene amplification, *MYC* enhancer amplification, and without any copy number change in either gene or enhancer. P-value of one-sided log-rank test is shown (n=100). **D)** Point mutations result in disruption of GATA2 binding sites which regulates *GFI1B* expression in AML. Shown tracks are histone modification ChIP-Seq data in human neurocrest cells and identified SNVs. **E)** Expression level of *GFI1B* in patients with and without enhancer mutations. P-value of one-sided t-test is shown (n=153). **F)** Kaplan-Meier plots of time to relapse for AML patients with and without *GFI1B* enhancer point mutations P-value of one-sided log-rank test is shown (n=153).

### Risk enhancer/promoter SNVs and small indels

We found that 346 patients in the TARGET cohort (69%) had SNVs/small indels located in enhancer/promoter regions that caused target expression change. For instance, we found 6 AML patients who have SNVs in the *GFI1B* +11k enhancer (30). The GATA2 binding sites in the enhancer were disrupted and *GFI1B* expression decreased correspondingly (Figure 3D, E). Growth factor independence 1b (*GFI1B*) encodes a key transcription factor regulating dormancy and proliferation of hematopoietic stem cells (HSCs) and the development of erythroid and megakaryocytic cells (31,32). Recent studies had revealed its critical role as a tumor suppressor in AML, as low *GFI1B* expression is associated with poor patient survival (33). Consistent with previous studies, the 6 patients with *GFI1B* enhancer mutation have significantly shorter time to relapse (Figure 3F). Another example involves 9 AML patients who have mutations in the *IDH2* +56k enhancer (Figure S11B). This enhancer is part of a known super enhancer of *IDH2*(34), and it is constitutively active in myeloid cell types. Previously, nonsynonymous mutations of *IDH2* have been reported in 9%-19% of adult AML patients, but relatively rare in childhood AML (<4%) (35,36). Mutations in *IDH2*, as well as in several other genes involved in regulation of DNA methylation (such as *DNMT3, TET2, IDH1*), were detected to have higher variant allele frequency (VAF) in adult AML patients, suggesting they were early mutational events in AML (37). We did not identify nonsynonymous *IDH2* mutations, but did find recurrent mutations in the *IDH2* enhancer. The mutations are predicted to disrupt ERG/FLI1 and E2F4 binding sites in the enhancer. The expression level of *IDH2* is significantly lower in patients with these mutations. Taken together, these results suggest a different mechanism of *IDH2* disruption in pediatric AML that was previously unappreciated. We also found 4 patients who have SNVs in the *GATA2* +126k enhancer and 2 patients who have SNVs in the *GATA2* promoter (Figure S11C). The enhancer was previously reported to be rearranged to vicinity of *EVI1* in adult AML patients (38). The mutations were predicted to disrupt FUBP1 binding site, and the expression of *GATA2* was significantly lower in the patients with mutations in enhancer/promoter regions.

### Coding and noncoding mutations affect distinct sets of genes and pathways

In total, our analysis of five pediatric cancer types have identified 1,175 genes recurrently altered in their coding regions, and 2,162 genes recurrently altered in their noncoding regions. Surprisingly, the overlap between the two groups of genes is very small (62 genes, 2%), suggesting general exclusivity of coding and noncoding mutations affecting a given gene (Figure 4A).

**Figure 4.**
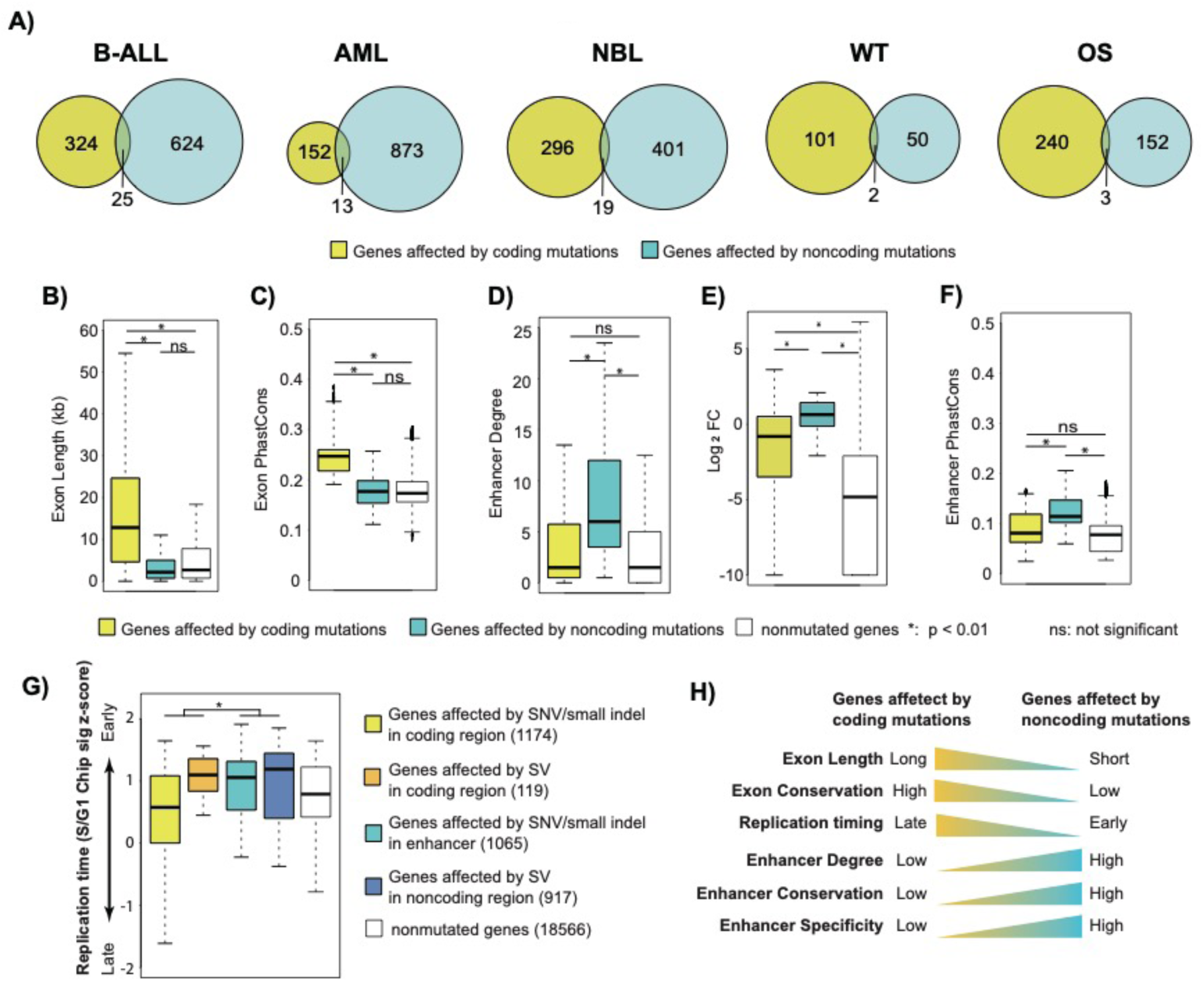
Coding and noncoding mutations affect distinct sets of genes. **A)** Venn diagrams of genes affected by recurrent coding and noncoding mutations in five cancer types. **B-G)** Features of genes affected by coding and noncoding mutations and genes without any mutation: gene exon length (B), exon conservation level measured by Phastcons score (C), enhancer degree (number of regulating enhancers) (D), gene expression specificity measured by fold change of average expression in a given cancer type compared to that of all five pediatric cancer types (E), enhancer conservation level measured by Phastcons score (F) and replication timing (G). **H)** Summary of different genomic features for the genes affected by coding and noncoding mutations. P-values of one-sided t-test are shown (n=21,841).

We investigated the genomic features of the genes affected by coding versus noncoding mutations. We found that genes affected by coding mutations are longer and have higher level of sequence conservation (Figure 4B, C). On the other hand, genes affected by noncoding mutations are linked to more regulating enhancers (Figure 4D), and their expression is more tissue-specific (Figure 4E). In addition, the regulating enhancers of those genes are more conserved (Figure 4F). Previous studies have suggested SNVs and small indels occur more frequently in genomic regions with late replication timing, while translocations and inversions occur more frequently in the genomic regions with early replication timing (39). Consistent with this trend, we found that genes affected by SNVs and small indels in coding regions are located in the relatively late replicating regions, while the genes in other groups have relatively earlier replication timing (Figure 4G). Pathway analysis reveals signaling pathways, including JAK/STAT, MAPK and Wnt signaling pathways, are mostly affected by coding mutations (Figure S12). In stark contrast, genes in metabolic pathways are mostly affected by noncoding mutations (Figure S12). Correspondingly, we found metabolic genes tend to be located in the early replicating region (Figure S13). Taken together, these data suggest the exclusivity of genes affected by coding versus noncoding mutations is likely due to the different genomic location and features of those genes (Figure 4H).

### Level of TF regulon disruption correlates with patient prognosis

Genes encoding lineage-specific transcription factors (TFs) are frequently mutated in pediatric cancers. For example, TFs that regulate B-cell development including *IKZF1, EBF1, PAX5, TCF3* are frequently altered in B-ALL patients (40-43). In addition, *MYCN* and *ZNF281* are known prognostic markers for neuroblastomas (44,45), while *RUNX1* and *CBFB* are also frequently altered in pediatric AML and are linked to unfavorable clinical outcomes (46).

In contrast to coding mutations of TFs, regulatory output of TFs can also be disrupted by mutations affecting individual enhancers/promoters/EP interactions regulating the target genes. Therefore, we compared the effects of coding and noncoding mutations on TF regulons (defined as the set of genes regulated by a TF) and patient outcome. For each cancer type, we report the top 10 most frequently affected regulons by combined coding and noncoding mutations. Most of the identified TFs have a known role in the specific cancer type (Figure 5A and Table S6). Furthermore, We found that regulon disruption by noncoding mutations occur more frequently than regulon disruption by coding mutations of the TF genes (Figure 5B). This is probably because mutations of a TF gene lead to a bigger disruption of the TF regulon, compared to mutations disrupting regulation of individual TF targets.

**Figure 5.**
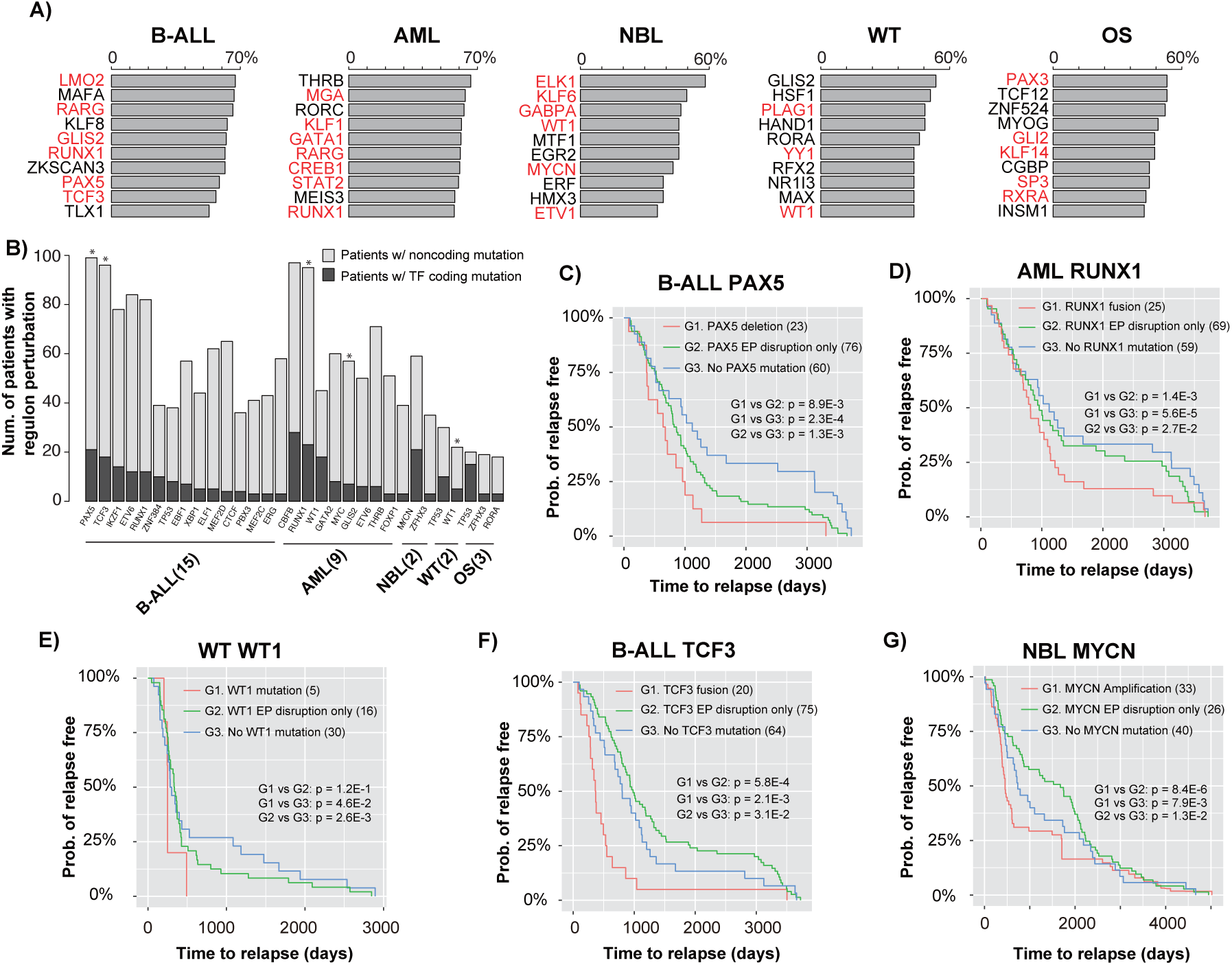
Degree of regulon disruption of key TFs is correlated with clinical outcome. **A)** Top 10 most frequently disrupted regulons in each cancer type. Bar plots show the percentage of patients with regulon disruption of the listed TFs; TFs with known role in the given cancer are highlighted in red. **B)** Number of patients affected by regulon disruption (either TF coding mutations or target gene noncoding mutations). Asterisks indicate TFs whose regulon disruptions are correlated with patient time to relapse (log-rank test p < 0.05, n=501). **C)** Kaplan-Meier plots of time to relapse for B-ALL patients with *PAX5* deletion, EP disruption involving PAX5, and without any mutation of the PAX5 regulon; **D)** Kaplan-Meier plots of time to relapse for AML patients with *RUNX1* fusion, EP disruption involving RUNX1, and without any mutation of the RUNX1 regulon. **E)** Kaplan-Meier plots of time to relapse for WT patients with *WT1* mutation, EP disruption involving WT1, and without any mutation of the WT1 regulon. **F)** Kaplan-Meier plots of time to relapse for B-ALL patients with *TCF3* fusion, EP disruption involving TCF3, and without any mutation of the TCF3 regulon; **G)** Kaplan-Meier plots of time to relapse for NBL patients with *MYC* amplification, EP disruption involving MYC, and without any mutation of the MYC regulon. P-values of one-sided log-rank test are shown. The numbers of samples are indicated in brackets.

We hypothesized that the level of regulon disruption is correlated with patient disease outcome. To test this hypothesis, we plotted time to relapse stratified by the degree of regulon disruption (*i*.*e*., TF coding mutation versus disruption of individual TF targets by noncoding mutations). Indeed, we found a strong correlation between time to relapse and the degree of regulon disruption. For instance, in B-ALL, patients with *PAX5* deletion have the shortest time to relapse, followed by patients with disruption of *PAX5* EP interactions. Finally, patients without any *PAX5* regulon mutation have the longest time to relapse (Figure 5C). We found the same correlations for *RUNX1* regulon in AML (Figure 5D), and WT1 regulon in WT (Figure 5E). For *PAX5, RUNX1*, and *WT1*, the mutations are all loss-of-function of the TFs. We also found correlations involving gain-of-function mutations of the TFs, including *TCF3* in B-ALL and *MYCN* in NBL. For TCF3, because the fusion event leads to gain-of-function of TCF3, patients with disruption of *TCF3* EP interactions have longer time to relapse compared to the patients without any TCF3 regulon mutation (Figure 5F). Presumably, the gain-of-function of TCF3 partially mitigate the effect of disruption of TCF target genes due to noncoding mutations. Similarly, NBL patients with disruption of *MYCN* EP interactions have longer time to relapse compared to the patients without any MYCN regulon mutation (Figure 5G). In summary, our data suggest that there is typically a wide spectrum of regulon disruption for key TFs in pediatric cancers. And the level of TF regulon disruption is associated with patient prognosis.

### Integrated mutation profiles suggest novel disease subtypes

Mutational profiles of coding regions have been widely used for patient stratification and prognosis purposes. This is not the case for noncoding mutational profiles. We therefore constructed mutational profiles of the five pediatric cancers using both coding and noncoding mutations. Clustering analysis of these combined mutational profiles enabled us to discover novel patient subtypes.

For B-ALL, 7 patient clusters are identified (Figure 6A). Of the 7 clusters, four are characterized by known translocation events in B-ALL including *ZNF384* rearrangement, *ETV6-RUNX1, TCF3-PBX1*, and *IGHV* translocation. All these are previously reported to be recurrent SVs in B-ALL (47,48). However, the mechanism by which these fusion proteins contribute to leukemogenesis has not been fully elucidated. Here, we found that in addition to creating fusion genes, the translocations can alter the expression of nearby genes by re-arranging locations of distal enhancers (Figure 6A, affected genes listed below heatmap).

**Figure 6.**
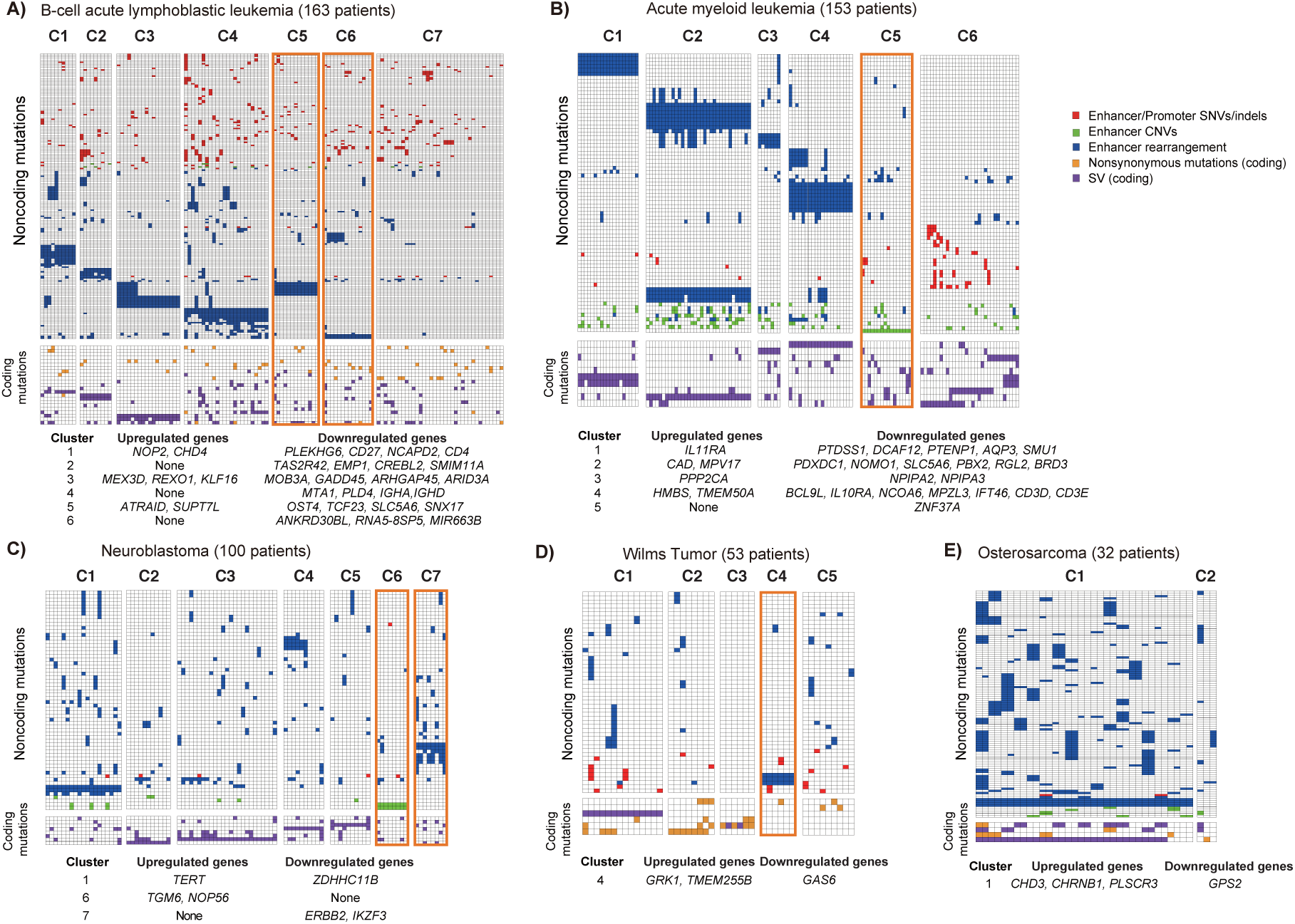
Integrative mutation signatures of pediatric cancers. **A)- E)** Mutation signatures of B-ALL, AML, NBL, WT, and OS patients. Heatmaps were generated using recurrently mutated genes (due to either coding or noncoding mutations) in at least 5 patients. Patients are clustered according to the combined coding and noncoding mutation profiles. The color of each cell indicates the class of mutations. Rows, genes. Columns, patients. For presentation purpose, the heatmaps are divided into submaps based on noncoding (top) or coding mutations (bottom). The tables below the heatmaps list up- or down-regulated genes due to noncoding mutations in all patients in a given cluster.

Two other clusters (C5 and C6) are characterized by novel inversion or translocation. Patients in C5 have Inv(2) (n=14). The inversion is predicted to hijack an enhancer to near *SUPT7L* and *ATRAID* and up-regulate the expression of these two genes (Figure S14A, B). *SUPT7L* is a subunit of the SPT3-TAFII31-GCN5L acetylase (STAGA) complex, which is known to regulate the stability of the *TCF3-PBX1* oncoprotein in ALL(49). Interestingly, we found Inv(2) significantly co-occurs with *TCF3-PBX1* in 6 patients (Figure S14C). Both *TCF3-PBX1* and Inv(2) are associated with aggressive clinical outcome in B-ALL. Patients who have both mutations show even shorter time to relapse compared to patients with either type of mutation (Figure S14D), suggesting synergy of these two pathways contributing to disease outcome.

Patient cluster C6 is characterized by the inter-chromosomal translocations at chr2 (t(2;7)(q21;q11), t(2;11)(q21;q11)). The translocations occur in 15 patients, and are predicted to disrupt the EP interaction involving the *ANKDR30BL* gene (Figure S15A), resulting in significant decrease of *ANKDR30BL* expression in patients with this translocation. Interestingly, *MIR663B*, a microRNA located in the intron of *ANKDR30BL*, is also down-regulated in those patients (Figure S15B). *MIR663B* is known to regulate the expression of *CCL17, CD40*, and *PIK3CD* in chronic lymphocytic leukemia (CLL) (50). Consistent with the previous finding, we observed significant expression increase of those genes in patients with *MIR663B* down-regulation (Figure S15B).

We also identified two potential novel cancer subtypes in NBL (C6 and C7, Figure 6C). Cluster 7 consists of 9 NBL patients with translocation on chr17 that results in enhancer rearrangement and down-regulation of *ERBB2* (t(1;17)(p33;q12), t(9;17)(p21;q12), t(11;17)(q13;q12)) (Figure S16A, B). *ERBB2* is essential for normal embryonic development and has a critical function in oncogenesis and progression of several cancer types including breast cancer, lung cancer, and leukemias (51-53). In addition, low *ERBB2* expression is associated with poor patient survival in NBL (54). Consistently, patients in cluster 7 have shorter time to relapse (Figure S16C). The other novel NBL subtype (C6) is characterized by amplification of the *TGM6* enhancer in 11 NBL patients and consequently increased *TGM6* expression in these patients (Figure S16D, E). Transglutaminase 6 (*TGM6*) is a protein associated with nervous system development (55). Transglutaminases, particularly *TGM2*, is known to play important roles in neurite outgrowth and modulation of neuronal cell survival (56). Our result suggests a potential oncogenic role of *TGM6* in NBL.

A novel AML subtype (cluster C5) is characterized by enhancer deletion of *ZNF37A* in 16 patients (Figure S17A). *ZNF37A* is involved in fusion events in breast cancer (57) and adult AML (58). Notably, AML patients with *ZNF37A* enhancer deletion have lower *ZNF37A* expression and shorter time to relapse (Figure S17B, C), suggesting the prognosis significance of this noncoding mutation.

A novel WT subtype (cluster C4) is characterized by enhancer rearrangement of *GAS6* in 6 patients (Figure S17D). Growth arrest specific 6 (*GAS6*) is a ligand for receptor tyrosine kinases *AXL, TYRO3* and *MER* whose signaling is implicated in cell growth and survival (59,60). Patients with the translocation have lower *GAS6* expression and shorter time to relapse (Fig S17E, F).

## Discussion

The landscape of noncoding mutations in pediatric cancers has not been comprehensively characterized. Here, we developed PANGEA, a general regression-based method to identify all classes of candidate risk noncoding mutations by joint analysis of patients’ mutations and gene expression profiles. Application of PANGEA led to a comprehensive and prioritized list of candidate risk noncoding mutations in five major pediatric cancers.

Previous studies on noncoding mutations have been focused on SNVs and small indels. In contrast, systemic analysis of SVs has been lacking. Due to their much larger sizes, SVs may have a bigger impact on shaping the cis-regulatory landscape than SNVs (61,62). In support of this notion, we indeed found that SVs are the most frequent class of risk noncoding mutations when adjusted for background occurrence frequency (Supplementary Fig S5). In total, our analysis has revealed 1,137 risk SVs affecting the expression of over 2,000 genes across five pediatric cancer types.

Previous studies have also been focused on fusion genes generated by SVs since they are intuitive candidates of driver events. However, only ∼30% of SVs generate fusion genes according to the current deposited SVs at ICGC. Moreover, ∼35% of the gene fusions in the TARGET cohort are generated via microhomology-mediated end jointing (MMEJ). Since microhomology tends to be located at the break points of nonpathogenic SVs (63,64), these data suggest the fusion genes may not be oncogenic drivers in cases of MMEJ. Instead, in our analysis, we found that 55% SVs altered regulatory landscape and expression of nearby genes of the break points. These genes warrant careful investigation for a role in oncogenesis in future studies.

We found that coding and noncoding mutations affect distinct sets of genes and pathways. This mutual exclusivity is likely due to the different genomic locations of these two classes of genes. We found that genes affected by noncoding mutations tend to be located in the region with early replication timing. This trend is consistent with previous reports that the occurrence of genomic rearrangements tends to occur in regions of early replication (65,66). Whether this correlation indicates a novel oncogenic mechanism needs to be further investigated. For instance, we found metabolic genes tend to be located in the early replicating region, and are more frequently affected by noncoding mutations. Rewiring of metabolism is a hallmark of cancer (67,68). Recent systematic analysis of TCGA data for 8 cancer types have reported that over 75% of metabolic genes are differentially expressed in each cancer type (69). However, it is unclear that to what degree the metabolism rewiring is mediated by noncoding mutations in those cancer types. Here our analysis suggests metabolic genes may be preferentially affected by noncoding mutation. In summary, our results highlight the need for comparative analysis of both coding and noncoding since novel cancer-related genes and pathways may be unveiled with comprehensive noncoding mutation analysis.

Many lineage-specific TFs are frequently perturbed in various cancers. However, the underlying oncogenic mechanism is challenging to understand. With comprehensive noncoding analysis, our approach provides a means to understand the detailed molecular mechanism underlying TF perturbation in cancers. For instance, recurrently de-regulated target genes of a TF suggest they can be important mediators of the disrupted TFs. Clinically, we found that the level of disruption of a transcription factor regulon can be used to stratify patient survival. More sophisticated computational models can be developed to prioritize target genes of perturbed TFs that contribute to oncogenesis.

## Methods

### Identification of single nucleotide variants (SNVs) and small indels

We used GATK (v3.8) and Freebayes (v1.0.2) (Table S9) to call SNVs and small indels. We first generated a set of initial SNV and indel calls with the default parameters of each software. Several filters were applied during the post-processing of the initial calls as suggested by the previous paper (70). First, we filtered mutations that overlap with low complexity regions. Second, we excluded regions with excessive read depth, as those regions are probably associated with spurious mappings. Third, we required the mutations to have multiple observations of the alternate (non-reference) allele in reads from both DNA strands. Last, we used the p-value cutoff of 0.01. SNVs passing these filters were intersected with annotations in dbSNP build 149. Calls matching both the position and allele of known dbSNP entries were removed (Figure S1A).

### Identification of structural variants (SVs)

Delly (v0.7.2) and Lumpy (v0.2.13) (Table S9) were used to call structural variants. We used the default parameter settings of both software with the exception of setting minimum mapping quality threshold to zero as advised by the Complete Genomics data analysis pipeline (https://target-data.nci.nih.gov/Public/Resources/WGS/CGI/READMEs/). The initial SVs called by both software were retained for further filtering. First, we removed SVs in which break points were located in repetitive regions. Second, we removed SVs that are also identified in the baseline genomes which consist of 261 WGS samples from the 1000 Genome Project (www.internationalgenome.org/data). Finally, we selected SVs with at least 7 supporting reads as the final set of SVs (Figure S2A).

### RNA-Seq data analysis

Raw reads from RNA-Seq were mapped to the reference human genome (release GRCh37) using STAR with default parameter setting. Transcripts were assembled using Cufflinks using mapped fragments outputted by STAR. Refseq (GRCh37) was used for the annotation of known transcripts. Normalized transcript abundance was computed using Cufflinks and expressed as FPKM (Fragments Per Kilobase of transcripts per Million mapped reads).

### Weighted elastic net model as a general framework for predicting risk mutations disrupting enhancer function

For a given gene promoter, we consider all types of mutations that could potentially disrupt its regulation, including SNVs and small indels, copy number variations, inversions, and translocations. These mutations could either disrupt the function of the cis-regulatory sequences *per se* or disrupt the interactions between the enhancer and the promoter. For the latter category of mutations, SVs could hijack enhancers for a given promoter. We define potential enhancer hijacking as existing enhancer relocated to new region with a nearby promoter (<200k bp). For each promoter, its potential enhancers are detected based on patient SV data. We developed a regression-based approach to identify specific mutations that are associated with gene expression change. We used elastic net to implement the regression analysis. Elastic net combines the strength of ridge regression and least absolute shrinkage and selection operator (LASSO). It can enforce sparsity, has no limitation on the number of selected variables, and encourages grouping effect in the presence of highly correlated predictors.

For each gene, let us consider a regression model with *n* observations (i.e. *n* patients). Suppose that *x*_*i*_ = (*x*_*ij*_, … *x*_*nj*_)^*T*^, *j* = 1, …, *p* are the predictors (mutations disrupting promoter regulation) and *y* = (*y*_1_, … *y*_*n*_)^*T*^ is the gene expression of the target promoters. *X* = [*x*_1_, …, *x*_*p*_] denotes the predictor matrix. The total number of predictors are the number of disrupting mutations summed over all promoters/enhancers regulating a given gene. The regression model can be expressed as *y* = *Xβ* + *ϵ* where *β* = (*β*, …, *β*_*p*_)^*T*^ and the noise term ε ∼ *N*(0, *σ*^2^*I*_*n*_). A model fitting procedure produces the estimate of *β*, 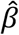. The elastic net model is defined as: 

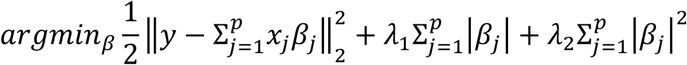

where *λ*_1_and *λ*_2_ are tuning parameters that balance the goodness-of-fit and complexity of the model.

Because of the enhancer-promoter links are computationally predicted by the IM-PET algorithm, we address the uncertainty of the prediction by proposing a weighted elastic net approach. Specifically, we associate the probability score of an enhancer-promoter prediction (computed by IM-PET) to the coefficient of the predictor in the model, to enforce penalty on predictors caused by uncertainty in EP predictions. The weighted elastic net model is as following: 

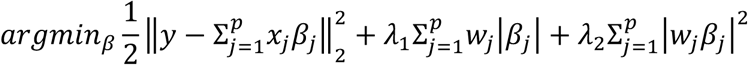

where *w*_*i*_ > 0, *j* = 1, …, *p* are weight based on the EP prediction score computed by IM-PET.

### CRISPR/Cas9-mediated translocation

sgRNAs targeting the break points were designed using CRISPOR (http://crispor.tefor.net) and cloned into the CRISPR vector pX459 (Addgene plasmid # 448139). CRISPR/Cas9-mediated translocation in Ba/F3 cells was performed as described in the previous study (71) with some modifications. Plasmids were co-transfected to Ba/F3 cells by electroporation. Dead cells were removed by centrifugation at 300x *g* and RT for 5 min 72 h post electroporation. Cell concentration and viability were measured using Countess II (Life Technologies). Live cells were re-suspended with Ba/F3 conditional medium to a concentration of 5 cells/mL. 100 μL of cell suspension was transferred to a 96-well plate and cultured for 2-3 weeks for selection of single cell clones. Genomic DNA was isolated using Quick-DNA 96 kit (ZYMO) and used for screening for clones with translocation by PCR (Table S7). Clones with translocation were further confirmed by Sanger sequencing.

### Generation of cell lines with stably expressed Cas9 endonuclease

The B-cell acute lymphoblastic leukemia cell lines NALM-6 and REH were lentivirally transduced with plasmid expressing Cas9 nuclease (LentiCRISPR V2; Addgene Plasmid #52961) as described below at a MOI of 0.5. Cells then underwent 14 days of antibiotic selection with media containing puromycin 0.25 ug/mL to generate cell lines that stably express Cas9. Both cell lines were cultured in RPMI media supplemented with 10% FBS and 100 unit/mL penicillin/streptomycin. All cell lines were validated by ATCC STR profiling and confirmed to be Mycoplasma free.

### sgRNA design

The sgRNAs targeting CHD4 were designed by Feng Zhang’s laboratory(72) (Table S8). The sequences for non-targeting control sgRNAs were chosen from previously described literature (73).

### Lentivirus production and transduction

Lentivirus was produced by transfecting HEK293FT cells with packaging plasmids pMD2.G and psPAX2, and the individual CRISPR components (Cas9 expressing plasmid or sgRNA expressing plasmid: MCB306, Addgene Plasmid #89360 or MCB320, Addgene Plasmid #89359). Lentivirus-laden supernatant was harvested at 48 and 72 hrs after transfection, and viral supernatant was filtered through 0.45 μm polyvinylidene difluoride filter (Millipore) and concentrated using ultracentrifugation at 25,000 rpm for 2 hrs at 4 °C. Virus was then tittered via transduction of target cell line (REH or NALM-6).

Cells were lentivirally transduced by spinfection with centrifugation of cells with virus in the presence of 8μg/ml polybrene (Millipore) at 1,000xg for two hrs. Wildtype cells were transduced with virus containing plasmid expressing Cas9 at a MOI of 0.5. Cells stably expressing Cas9 were then transduced with plasmid containing sgRNA targeting CHD4 or non-targeting control at a MOI of 0.4. Three days after virus transduction, GFP-positive (transduced) cells were sorted by FACS, then seeded into 6-well plates for recovery before the following analyses.

### Data and Software availability

Data and software used in this study are noted in the Method Details section above and Table S9. In this study, we developed a new software, PANGEA which is deposited at https://github.com/tanlabcode/PANGEA.

## Acknowledgments

This work was supported by National Institutes of Health grants GM104369, GM108716, HG006130, CA226187, CA233285 (to KT), K08CA184418 (to SKT), and a pilot grant from the Children’s Hospital of Philadelphia Research Institute (to KT and SKT).

## Author contributions

B.H. and K.T. designed the overall study, analyzed data, and wrote paper. P.G. performed most of the experiments. C.H.C and Y.D. performed competition growth assays. C.C., Y.D, G.C., H.K, S.K.T, S.P.H helped with experimental design, data interpretation, and manuscript writing.

